# Guidestar: a spike-in approach to improve RNA detection accuracy in imaging-based spatial transcriptomics

**DOI:** 10.1101/2025.01.09.632272

**Authors:** Jazlynn Xiu Min Tan, Lurong Wang, Wan Yi Seow, Jeeranan Boonruangkan, Mike Huang, Jiamin Toh, Kok Hao Chen, Nigel Chou

## Abstract

Imaging-based spatial transcriptomics technologies, such as MERFISH and seq-FISH, use combinatorial barcoding and imaging to simultaneously detect individual RNA molecules from 10s to 10,000s of genes. These technologies require the decoding of individual RNA molecules’ location and gene identity from stacks of images. However, beyond using ‘blank’ code-words as negative controls, there is a lack of ground truth information embedded within the assay to experimentally measure the accuracy and sensitivity of the decoding algorithm. We introduce Guidestar, a system of spike-in controls integrated within a combinatorial FISH assay, that labels a subset of RNA transcripts with additional probes. These probes are imaged separately as ‘guide bits’, which serve as ground-truth data to assess decoding accuracy at the level of individual RNA molecules. Using Guidestar to evaluate accuracy of an existing decoding method suggested alternative parameter settings that increased sensitivity with minimal impact on accuracy. We also used the Guidestar dataset to train a machine-learning based classifier to distinguish true from false RNA calls, yielding 9% and 40% higher F1 scores across cell line and tissue samples, respectively.

## Introduction

Spatial transcriptomics technologies reveal gene-expression profiles of cells together with their spatial context in a tissue, promising novel insight into tissue function and disease etiology. A major subset are multiplexed fluorescence in situ hybridization (mFISH) based approaches, including MERFISH^1^, seqFISH^2,3^ and a variety of commercially available options (10x Xenium, Vizgen MERSCOPE, Nanostring CosMX etc.), that simultaneously measure 100s to 10,000s of individual RNA molecules using oligonucleotide probes tagged with fluorescent dyes. Each RNA species is assigned a different set of read-out probes corresponding to a binary code-word, and read out over multiple imaging cycles. Hence, RNA location and identity must be decoded from stacks of noisy images, a core analysis step preceding all downstream analysis, including cell segmentation^4^, cell type clustering^5–8^, spatial domain segmentation^9–12^, identification of spatially varying genes^13,14^ and cell-cell interaction analysis^15,16^.

Current approaches begin decoding either by assigning a RNA identity to each pixel and grouping contiguous, similarly assigned pixels into RNA spots^17–19^, or by matching detected spots across bits^1–3^. The former approach is generally more computationally efficient and is the focus of this study. Subsequently, the identified RNAs are conventionally filtered based on 3 features, namely spot size, mean intensity, and Hamming distance to the corresponding RNA barcodes. The filtering step aims to improve accuracy by eliminating falsely detected RNA calls, but may also remove substantial numbers of true RNA calls, leading to decreased sensitivity.

The output of a multiplexed FISH assay is most commonly evaluated by comparing inferred cell type abundances with scRNA-seq^20,21^, but this confounds RNA decoding accuracy with that of subsequent analysis steps including cell segmentation and cell type clustering. The quality of RNA decoding can be more directly validated by comparison to bulk RNA-sequencing data, which correlates the relative abundances of different RNA species at the whole-sample (bulk) level but is unable to assess decoding and spatial accuracy at the level of individual genes. Therefore, there is a critical need for more direct methods to assess sensitivity and accuracy in current FISH-based spatial transcriptomics workflows.

Hence, we developed Guidestar, a spike-in control approach that labels a subset of RNA transcripts with an additional set of probes (**Figure 1a**). These probes are imaged as part of the MERFISH assay but on separate hybridization cycles: this approach allows us to evaluate if the spots detected in the “guide” images spatially colocalize with the decoded MERFISH spots, allowing for validation of RNA decoding at the single-molecule level. Therefore, they serve as a matching ground truth dataset to evaluate detection accuracy for this subset of RNA transcripts. Importantly, the “guide” probes are integrated with the MERFISH probe-library through the same probe-design process and introduced with MERFISH encoding probes in a single hybridization step.

**Figure 1:**
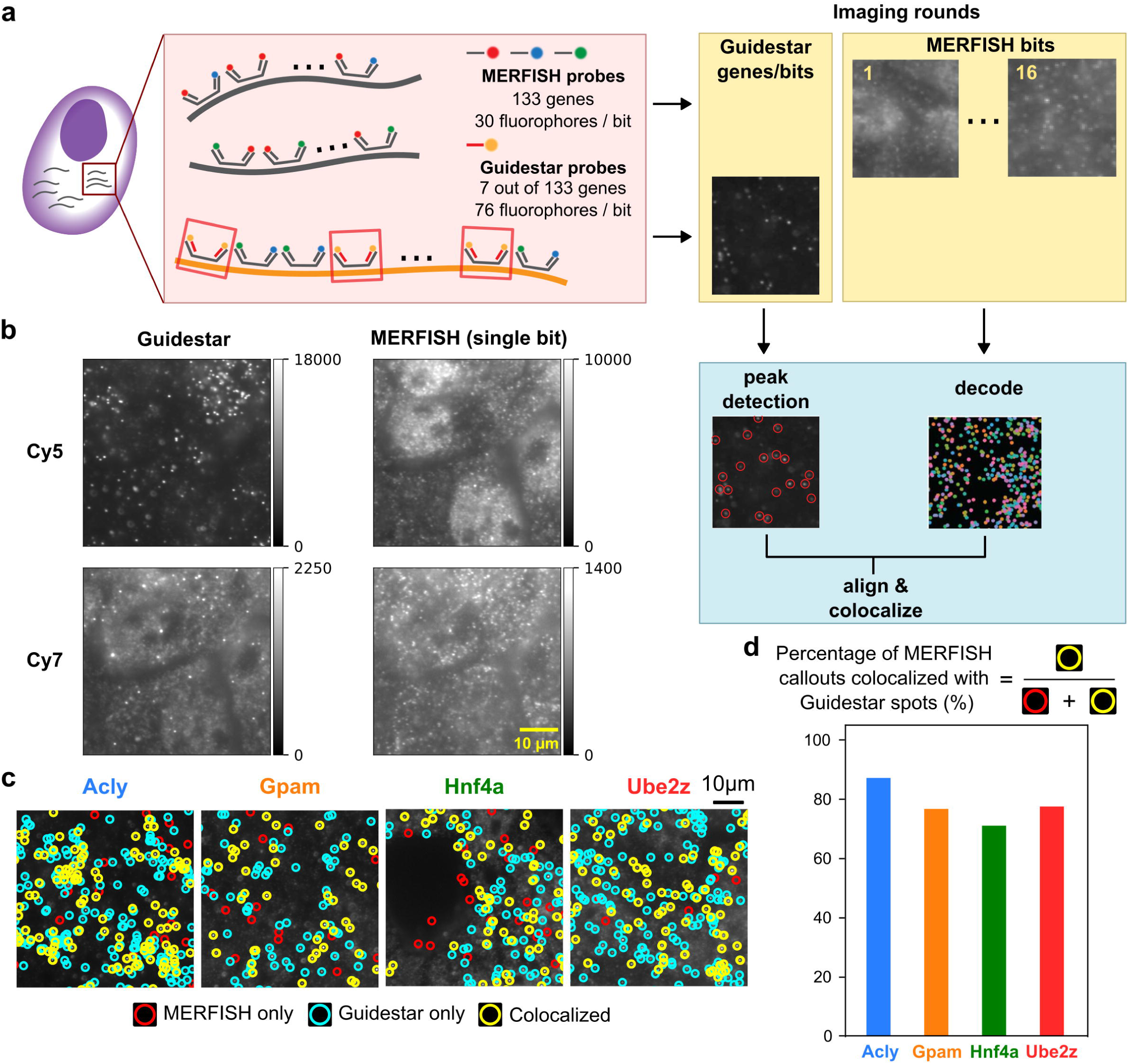
Guidestar assay. **(a)** Schematic showing Guidestar (GS) probe design and imaging protocol. GS probes are interspersed with MERFISH probes on a small subset of genes from the MERFISH library. GS uses 2.5 times the number of fluorophores compared to a single imaging round in MERFISH. GS images are acquired, one gene at a time, before performing the combinatorial MERFISH assay. GS RNA spots are identified by peak finding using local intensity maxima, while MERFISH images are decoded by matching voxel sequences to the codebook. The spots identified in GS are colocalized with those decoded via MERFISH. **(b)** Representative raw images (display range from 0 to 99.9% percentile of the pixel intensities) from two color channels for GS and MERFISH. **(c)** Example GS images overlaid with the locations of identified GS peaks and MERFISH callouts. Red circles: spots called out by MERFISH only, yellow circles: spots detected by GS peak-detection only, blue circles: spots detected by both MERFISH and GS. **(d)** The percentage of MERFISH merfish spots colocalized with GS spots for each gene.

## Results

We designed an integrated library that combined a 133-gene MERFISH panel containing genes of interest in liver tissue with Guidestar probes targeting 7 of the 133 genes. The subset of genes targeted by Guidestar probes were chosen to have (1) sufficiently long transcripts to accommodate the additional probes and (2) moderate to high expression levels in the tissue of interest as estimated from bulk measurements to ensure sufficient numbers of detected RNA spots. We then hybridized this oligonucleotide probe-set with samples from the AML-12 murine hepatocyte cell line and performed the Guidestar-integrated MERFISH assay protocol (**Figure 1a**). After performing quality control checks on the 7 Guidestar genes and removing genes with low FPKM on the cell line sample (**Figure S1a**), we selected 4 Guidestar genes (*Acly*, *Gpam*, *Hnf4a* and *Ube2z*) for further analysis.

We observed that the Guidestar images had superior image quality relative to MERFISH images, with RNA spots that could be clearly distinguished from background fluorescence (**Figure 1b**). To verify the improvement in Guidestar image quality over MERFISH, we quantified the drop off in intensity from the center of each spot to the periphery (Methods section *Signal and background brightness comparison*, **Figure S1b**), showing that Guidestar spots had at least 2-fold improved signal-to-background ratio (SNR) at the spot center relative to spots in the MERFISH imaging rounds.

We then assessed the extent of colocalization between Guidestar and MERFISH spot locations, defining colocalization as a maximum euclidean distance of 0.48 μm (4 pixels) based on percentages of colocalization at different distances (**Figure S2a**). For each of the Guidestar imaging rounds (corresponding to individual chosen genes), we identified RNA spot locations by detecting local maxima^22^. MERFISH images were analyzed (preprocessing, registration and decoding) as previously described to obtain RNA identities and locations^1,23^. We found that across the 4 Guidestar genes, 71 - 87% of spots called by MERFISH colocalized with Guidestar (**Figure 1c, d, Figure S2b**). In comparison, colocalization percentages between MERFISH and unmatched guidestar genes (e.g. Acly Guidestar spots vs Gpam MERFISH spots), which represent negative controls, had a mean colocalization of 4.0% across the 12 gene pairs (**Figure S2b**). On the other hand, 27 - 44% of spots detected by Guidestar were also detected by MERFISH (**Figure S2e**). The fraction of spots detected by both Guidestar and MERFISH out of all Guidestar spots was similar to the expected detection efficiency of MERFISH in high-abundance libraries of unexpanded cells (MERFISH detected 21±4% of RNA spots determined by smFISH previously)^24^. Together, these tests suggest that Guidestar can be used as a ground-truth reference to evaluate MERFISH RNA decoding results.

### Guidestar enables parameter tuning of existing MERFISH decoding pipelines

A key step in MERFISH decoding is the use of the number of detected negative controls (specifically “blank” codewords) to tune the stringency of RNA callout filtering^4,18^. “Blanks” are codewords that do not encode any gene in the library but are also at a minimum Hamming distance of 4 away from the gene-encoding codewords, and hence serve as a negative control. For this experiment, 7 out of the 140 MERFISH codewords were assigned to blanks. The hyperparameter “target misidentification rate” measures the fraction of “blank” codewords that would be called out as true spots after filtering. This is often set heuristically at 0.05 (i.e. 5% of the “blanks” would be accepted as true RNA) during filtering^19^. We will refer to this method of RNA spot filtering as the blank fraction threshold (BFT) approach.

Since our Guidestar dataset uniquely provides ground truth of the true positive RNA callouts, we used Guidestar to estimate the precision and recall of the BFT approach (**Figure 2a**), and varied the misidentification rate to determine the optimal threshold that balances precision and recall, using the F1 score as a summary statistic. We found that at the default 0.05 misidentification rate, BFT has high precision but low recall, suggesting that the BFT approach at default settings may be biased toward removing false positives, while also filtering out a large proportion of true spots. Our analysis in this cell line sample shows that a more optimal trade-off in precision and recall could be achieved at a higher misidentification rate of 0.2-0.4 (**Figure 2b**), which maximizes the F1 score for the Guidestar genes. These parameter settings yield a substantially higher sensitivity, with 23% - 53% more RNA callouts relative to default settings. This analysis illustrates the value of Guidestar for quantifying both the sensitivity and accuracy of decoding methods, and its utility for optimizing hyperparameters for decoding.

**Figure 2:**
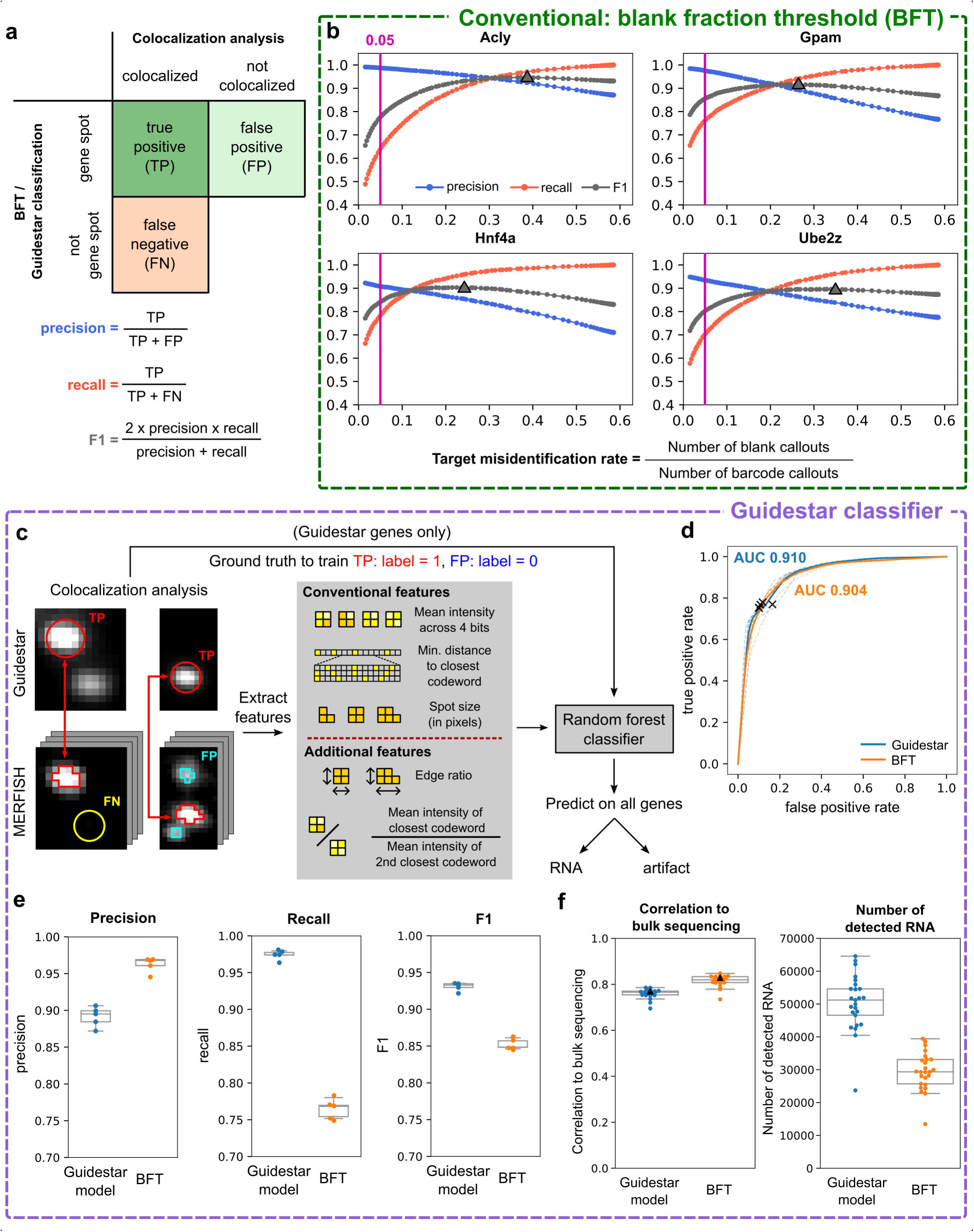
Application of Guidestar to cell line data. **(a)** Colocalization of MERFISH callouts with Guidestar (GS) spots (demonstrated in **c**) was used to designate callouts as true positives, false positives and false negatives for calculation of precision, recall and F1. **(b)** Precision, recall and F1 plotted as a function of the target misidentification rate, a parameter in the conventional decoding pipeline (representing the fraction of barcode callouts that correspond to blank controls after filtering). Triangle marker denotes maximum F1 score. **(c)** Spot features are extracted from true positive, false positive and blank MERFISH callouts of the training data consisting of GS genes to train the random forest classifier. The model is then applied to the rest of the gene callouts to classify the callouts as RNA or artifact. **(d)** GS model and BFT receiver operating characteristic curves (dashed lines: curves for each FOV in test set, solid lines: average across all FOVs). BFT performance at 0.05 misidentification rate is indicated by the crosses for each FOV. Area under the curve (AUC) values are indicated for the average ROC. **(e)** Precision, Recall and F1 for the GS model and BFT (at default 0.05 misidentification rate). Center line, median; height of the box, interquartile range (IQR); whiskers, 1.5 × IQR. Points represent individual FOVs in the test set. **(f)** Trained GS model and BFT applied to the full gene library. Log correlation to bulk RNA-sequencing fragments per kilobase million (FPKM) values (left) and total number of detected RNA molecules (right). Center line, median; height of the box, interquartile range (IQR); whiskers, 1.5 × IQR. Points represent individual FOVs within the dataset.

### Guidestar enables the design of binary classifiers for spot filtering

We then sought to use the Guidestar-derived ground-truth to train a binary classifier (Random Forest [RF] model) for MERFISH spot filtering (**Figure 2c**). We first set aside 5 out of 25 Fields of View (FOVs) as the test set. We performed 5-fold cross validation to optimize the number of genes used (**Figure S3a,** Methods section *Quality control and Guidestar gene selection*). All 4 genes (*Acly, Gpam, Hnf4a* and *Ube2z*) were used as the number of genes did not have a significant effect. As the number of positive examples (colocalized) and negative examples (MERFISH only spots) differed, we also optimized class balancing (Methods section *Class balancing approach*) with a combination of downsampling and augmentation with ‘blank’ codewords (**Figure S3c**). We expanded the features used for classification from the 3 conventional parameters: mean intensity of pixels within the spot, the hamming distance between the imaged codeword and closest theoretical barcode, and the spot size, to include an additional 12 intensity, shape, and distance-related features (**Table S4**), then used sequential forward feature selection to obtain the reduced set of 4 features (Methods section *Additional features and feature selection*) that produced the best F1 score during cross validation (**Figure S3e**). Finally, we conducted hyperparameter optimization for the RF classifier (**Figure S3f**), and selected the model with the best mean F1 score across 5 folds. We then retrained the model on the full training set and evaluated its performance on the held-out test set.

We first compared the performance of the classifier (methods *model training*) with that of the BFT approach at default settings (0.05 misidentification rate). On the test set, using different FOVs as technical replicates, the RF model obtained a mean precision of 0.89 and recall of 0.97 while BFT (default settings) had a mean precision of 0.96 and recall of 0.76. The RF classifier had a significantly higher F1 score compared to BFT (0.93 for RF vs 0.85 for BFT; *p* = 3.68 x 10^-5^, two-tailed paired student’s t-test, **Figure 2e**). To comprehensively compare the performance of the classifier and BFT, we computed the receiver operating characteristic (ROC) curves for both and found that they had similar area under curve (AUC), with the classifier being marginally higher (0.910 for classifier, 0.904 for BFT; *p* = 2.16 x 10^-5^, two-tailed paired student’s t-test) (**Figure 2d**). This suggests that the performance of BFT is comparable to the classifier across misidentification rate settings. However, the classifier at the default output probability threshold (0.5) generates a higher F1 than the BFT at its default setting of 0.05 misidentification rate.

Since Guidestar yielded improved decoding results in the AML-12 cell line sample, we reasoned that it would be valuable for spot filtering in tissue samples, where the presence of extracellular matrix and more densely packed cells often result in higher background autofluorescence and nonspecific probe binding, leading to more challenging decoding relative to cell line samples. Using the same procedure described above for cell line samples, we applied our Guidestar integrated probe library to a mouse liver tissue sample. We selected *Acly*, *Gpam*, *Hnf4a*, and *Pigr* as Guidestars after quality control (**Figure S1a**).

Compared to the cell line dataset, there were fewer positive examples relative to blanks or MERFISH-only spots due to higher RNA densities and smaller cell size in tissue. The percentage of MERFISH callouts that were colocalized to Guidestar ranged from 14% to 29%. In contrast, 4.3% of unmatched genes colocalized (**Figure S2c, d**). As there were more MERFISH-only spots than colocalized spots, we downsampled the negative examples (MERFISH-only spots) to match the number of colocalized spots to avoid class imbalance (**Figure S3d**). We then trained the classifier (using the optimized hyperparameters determined from the cell line sample) on the liver tissue dataset (training set = 40 FOVs, held out test set = 9 FOVs).

On the test set, using different FOVs as technical replicates, the RF model obtained a mean precision of 0.58, recall of 0.79, and F1 of 0.65 (**Figure S4a**). This was a significant improvement over the default BFT method which, despite a higher mean precision of 0.67, had a lower recall of 0.39 (49% that of the RF model) and a significantly lower F1 of 0.46 (p = 2.99 x 10^-3^, two-tailed paired student’s t-test). Notably, Guidestar achieved a larger improvement of 41.3% in F1 score for the tissue sample compared to 9.22% on the cell sample. We again computed the ROC curves for both methods. Here, the classifier had a similar AUC to BFT (0.830 vs 0.807; p = 0.111, two-tailed paired student’s t-test, **Figure S4b**).

Next, we tested the Guidestar-trained classfier’s ability to generalize to unseen RNA species on both cell line and tissue samples. We applied the trained model to the full probe library where the majority of genes were not imaged with corresponding Guidestar probes. For the cell line sample, the RF model resulted in a small but significant decrease in mean FPKM correlation across all FOVs (0.76 for RF model vs 0.82 for BFT; p = 2.26 x 10^-22^, two-tailed paired student’s t-test), but also yielded on average 71.4% higher spot callouts per FOV than BFT (**Figure 2f**). In the tissue sample, we also observed a decrease in mean FPKM correlation across all FOVs (0.58 for RF model vs 0.64 using BFT, p = 3.33 x 10^-11^, two-tailed paired student’s t-test). However, the RF model trained with Guidestar called out 3.1-fold more spots on average per FOV compared to BFT (**Figure S4c**). This is consistent with our validation results above on the guidestar genes, which showed that the RF classifier is less precise but yielded a substantial increase in sensitivity. These results suggest that our model, trained on a subset of genes, can be generalized to unseen genes.

## Discussion

Localizing and decoding RNA species from raw images is a critical step for analyzing in-situ high-throughput transcriptomic data. However, methods for systematically validating this analysis step have been lacking. To address this gap, we have developed Guidestar, a method designed around spike-in ground truth controls in MERFISH assays, enabling the validation and optimization of existing RNA decoding methodologies. We demonstrated that Guidestar probes, when applied to a small subset of genes, yield high-quality images with clearly distinguishable RNA spots, and that a significant proportion of these spots colocalize with decoded MERFISH callouts. Importantly, despite imaging only a small subset of genes with Guidestar probes, classifiers trained on this data generalized well to the remaining genes in the library due to the use of common spot features that are not gene specific.

The primary advantage of Guidestar is its ability to directly validate both the precision and sensitivity of existing MERFISH decoding approaches. We evaluated the common approach of utilizing the BFT to filter out false spots. Our sensitivity and precision analyses on a cell line dataset indicate that the default setting for BFT (0.05 misidentification rate) may lead to the removal of a substantial fraction of true spots, with only marginal improvements in accuracy relative to slightly higher settings for this parameter. This underscores the value of Guidestar in assessing the trade-off between precision and recall, allowing users to fine-tune decoding settings, whether prioritizing conservative results with lower RNA count or maximizing RNA detection with slightly elevated error rates. Moreover, in the presence of sample-specific image-quality fluctuations driven by tissue-specific fluorescent background and varying probe permeability and binding, Guidestar enables the important task of spike-in based parameter optimization. Thus, we believe that Guidestar holds considerable promise for improving the reliability of in-situ RNA imaging across diverse tissue types.

Furthermore, our study highlights the limitations of relying solely on the correlation to bulk FPKM values as a metric. The FPKM correlation metric is highly sensitive to precision, since incorrect RNA calls skew the relative abundances of detected genes. However, it does not account for detection sensitivity, since correlation values remain high even if only a small fraction of total detectable RNA molecules are called by the decoder. This may lead to biased decoding favoring precision over recall. To address this limitation, there is a critical need to supplement FPKM correlation with methods that estimate detection sensitivity. To our knowledge, Guidestar is the only method that provides a true positive reference from the same tissue slice, enabling accurate estimation of both recall and precision.

Beyond validation of existing methods, the ground truth reference produced by Guidestar can also be leveraged for developing improved decoding approaches, such as the classifier we developed as a proof of concept. Our results demonstrate that the Guidestar-trained classifier significantly improved the F1 score compared to the BFT approach at default parameters, across both cell line and tissue samples. However, the ROC curves, while slightly higher for the classifier, revealed comparable performance between the two methods, underscoring the robustness of current decoding methods and importance of parameter tuning for optimal decoding outcomes. Notably, the trained models generalized to non-Guidestar genes, suggesting the potential of Guidestar for developing and testing more sophisticated spot detection approaches. The classifier used in this study is limited by its reliance on existing decoding pipelines that first identify candidate RNA spots which can then be classified as true RNA spots or false positives. A promising future direction would be to apply more sophisticated models to decode RNA spots directly from raw image stacks, such as with semi-supervised deep-learning^25^, or by adapting deep-learning models used for smFISH spot detection^26^. In both cases, using Guidestar as ground truth for training or model refinement could greatly enhance accuracy while reducing the reliance on labor-intensive and subjective manual annotation.

In summary, Guidestar is a conceptually innovative technique that combines spike-in methodology, commonly employed in RNA-seq^27^ and scRNA-seq^28^, with machine-learning approaches to validate MERFISH RNA decoding results directly. We envision Guidestar being utilized as a quality control and validation tool in multiplexed FISH workflows, enhancing the quality of downstream results derived from spatial omics assays.

### Limitations of Study

The accuracy of ground truth spot coordinates derived from Guidestar images may be influenced by raw image quality and the employed peak detection algorithms. Suboptimal assay conditions, relating to instrumentation, probe design and hybridization, can compromise image quality, leading to reduced accuracy of Guidestar and MERFISH spot calls. Peak calling in Guidestar images, similar to smFISH, is sensitive to algorithm choice and peak height thresholds. Recent advancements in deep learning for spot detection^25,26^ may offer potential improvements in accuracy and automation.

The classifier model presented here serves as a filter for existing spots that operates in the context of a larger analysis pipeline preceded by a spot-calling step, rather than a comprehensive method to decode RNA spots from the raw data. The model also does not leverage information from the Guidestar spots that do not colocalize with MERFISH. These spots could correspond to additional true RNAs undetected by current MERFISH labeling, and may represent potential false negatives to augment the training examples used in the current model.

## Resource availability

### Lead contact

Further information should be directed to the lead contact, Nigel Chou.

### Materials availability

This study did not generate new unique reagents or materials. A list of reagents used and their catalog numbers can be found in the key resources table.

### Data and code availability

● Raw images can be found on Zenodo (will be made available upon publication).
● Training and MERFISH data can be downloaded from https://drive.google.com/drive/folders/122UvRUf9SqZhPW2tYLKfgjZl1Y_HP1Kx?usp=sharing
● The Guidestar code can be downloaded at https://github.com/nigelchou/Guidestar DOIs are listed in the key resources table.

## Supplemental information

**Document S1.** Figures S1-S5. Tables S2-S4.

**Document S2.** Table S1.

## Acknowledgements

This work was funded by Agency for Science, Technology and Research (A*STAR) grant no. I1801E0029 (K.H.C, J.B., M.H.), the Singapore Ministry of Health’s National Medical Research Council (NMRC) grants OFIRG20nov-0056 (K.H.C., W.Y.S.) and OFYIRG23jul-0031 (N.C, W.L.), the National Research Foundation Competitive Research Programme (CRP) grant no. NRF-CRP25-2020-0001 (K.H.C., J.B., W.L.), and A*STAR IAF-PP-H18/01/a0/020 (K.H.C., JM.T.). J.T. and N.C. were also supported by the A*STAR Graduate Academy. The funders had no role in study design, data collection and analysis, decision to publish or preparation of the manuscript. We thank Shyam Prabhakhar and Lee Hwee Kuan for useful discussions and Chung Pui Paul Cheng for assistance in probe library preparation.

## Author contributions

Conceptualization: K.H.C., M.H., J.T., N.C.; Data Curation: J.T., J.B.; Formal Analysis: J.T., L.W., J.B., W.Y.S., N.C., K.H.C.; Funding Acquisition: K.H.C., N.C.; Investigation: W.Y.S., JM.T. ; Methodology: J.T., J.B., M.H., N.C., K.H.C.; Resources: N.C., K.H.C.; Software: J.T., L.W., J.B, M.H., N.C.; Supervision: K.H.C., N.C.; Validation: W.Y.S., L.W., J.T.; Visualization: J.T., L.W.; Writing—original draft: N.C., K.H.C., J.T, W.Y.S.; Writing—review & editing: N.C., J.T., K.H.C. W.Y.S.

## STAR Methods

### Probe Design

30-nt targeting regions for the Guidestar and MERFISH probes were identified using a previously published algorithm^23^. Transcript sequences from the GENCODE website (mouse m4) were used as reference. A specificity table was calculated using 15-nt seed and 0.3 specificity cut-off was used for MERFISH probes, while a 0.5 specificity cut-off was used for Guidestar probes. Quartet repeats (’AAAA’, ’TTTT’, ’CCCC’, and ’GGGG’) were excluded from the possible target regions. We randomly distributed the Guidestar and MERFISH probes on the target regions. The list of encoding probes can be found in **Table S1**, the list of readout probes can be found in **Table S2**, and the codebook can be found in **Table S3**.

### Probe library amplification and preparation

The probe library (Twist Bioscience) was amplified following a previously published protocol^1^. First, we amplified the oligonucleotide pool by limited-cycle PCR using Phusion Hot Start Flex 2X Master Mix (NEB, cat. no. M0536L), using an annealing temperature of 66°C. The T7 promoter sequence was introduced on the reverse primer during PCR. Then, overnight in vitro transcription (IVT) was done using a high-yield IVT kit (NEB, cat. no. E2050S), before reverse transcription was done on the RNA template (Thermo Fisher, cat. no. EP0753). The RNA was then cleaved by alkaline hydrolysis, generating single-stranded DNA (ssDNA). Finally, the ssDNA were purified using magnetic beads (Beckman Coulter, cat. no. A63882) and eluted with nuclease-free water. The probes were freeze dried and stored at −20°C until use. The following primers were used for PCR: 5′-TGGTTCAATCGTATGCCCGT-3′ and 5′-TAATACGACTCACTATAGGGGTCACTTAGCCAACGCCGAT-3′.

### Preparation of cell-culture samples

We cultured AML12 (ATCC CRL-2254) cells in high-glucose DMEM (Hyclone, cat. no. SH30022.01) that was supplemented with 10% FBS (Thermo Fisher, cat. no. 26140079). The cells were grown to ∼80% confluency on autoclaved 40 mm round coverslips (Warner Instruments, cat. no. 64–1500) placed in 60 mm dishes. The samples were then fixed in 4% vol/vol paraformaldehyde (Electron Microscopy Sciences, cat. no. 15714) in 1× PBS for 15 min at room temperature. After fixation, the cell samples were quenched with 0.1 M glycine (1st BASE, cat. no. BIO-2085) for 1 min at room temperature before rinsing with 1× PBS. Fixed samples were stored in -80°C until use. Prior to usage, samples were permeabilized in 70% ethanol overnight at 4°C.

### Coverslip functionalization

Coverslips were functionalized prior to tissue sectioning^29^. The coverslips (Warner Instruments, cat. no. 64–1500) were cleaned by sonication in 1M KOH using an ultrasonic water bath for 20 min, then rinsed with MilliQ water thrice followed by a rinse with 100% methanol. Next, the coverslips were immersed in an amino-silane solution (3% vol/vol (3-aminopropyl)triethoxysilane (Merck cat no. 440140-500ML), 5% vol/vol acetic acid (Sigma, cat. no. 537020) in methanol) for 2 min at room temperature before being rinsed thrice with MilliQ water. The functionalized coverslips were then air-dried overnight and used immediately or stored in a dry environment at room temperature for up to two months.

### Tissue sample preparation

All animal care and experiments were carried out in accordance with Agency for Science, Technology and Research (A*STAR) Institutional Animal Care and Use Committee (IACUC) guidelines (IACUC #211580). Histology work was performed by the Advanced Molecular Pathology Laboratory, IMCB, A*STAR, Singapore. Briefly, 8 week old C57BL/6NTac female mice (InVivos) were euthanized with ketamine, and their livers were collected. The livers were cut into smaller pieces and frozen as soon as possible in Optimal Cutting Temperature compound (Tissue-Tek O.C.T.; VWR, cat. no. 25608–930). The tissue blocks were stored at −80°C. 7 μm sections of the tissue blocks cut with a cryotome onto the functionalized coverslips. After air drying for 5-10 min at room temperature, samples were fixed in a 4% vol/vol paraformaldehyde in 1× PBS solution for 15 min. Samples were then rinsed with 1× PBS and stored at −80°C for future use. Samples were permeabilized in 70% ethanol overnight at 4°C before use.

### Guidestar and Multiplexed FISH experiments

After permeabilization, the samples were rehydrated in 2× saline–sodium citrate (SSC, Axil Scientific, cat. no. BUF-3050-20×1L) for 5 min. Samples were then pre-hybridized in a 20% formamide wash buffer, containing 20% deionized formamide (Ambion, cat. no. AM9342, AM9344) and 2× SSC, at 47°C. AML-12 cell samples were pre-hybridized for 30 min, while the mouse liver section samples were pre-hybridized for at least 3 hours. The library probes were diluted to a concentration of 100 μM in 20% hybridization buffer that consists of 20% deionized formamide (vol/vol), 1 mg/ml yeast tRNA (Invitrogen, cat. no. 15401011) and 10% dextran sulfate (Abcam, cat. no. ab146569) (wt/vol) in 2× SSC. The samples were stained with the encoding probes at 47°C overnight in a humidified, air-tight container. We then washed the samples in a 20% formamide wash buffer at 47°C for 15 min, twice. Finally, we washed the samples twice with 2× SSC before being imaged immediately or stored for no longer than 24 h at 4°C in 2× SSC before imaging.

### Guidestar and Multiplexed FISH imaging cycle

All datasets were acquired on a homebuilt imaging system described previously^23^. The setup was based around a Nikon Ti2-E body, an Andor Sona 4.2B-11 sCMOS camera. We used a Nikon CFI Plan Apo Lambda ×60 1.4-n.a. oil-immersion objective for all imaging experiments. For illumination we used MPB Communications fiber lasers for Cy5 (647 nm) and IRDye 800CW (750 nm), respectively: 2RU-VFL-P-1000-647-B1R (1000 mW) and 2RU-VFL-P-500-750-B1R (500 mW). Samples were secured to the microscope stage using the Bioptechs FCS2 flow chamber, and a custom-built computer controlled fluidics system was used to perform sequential rounds of readout hybridization and imaging. For each round of imaging, readout probe solution was flowed into the chamber and incubated with the sample for 30 mins at room temperature. The readout probe solution consisted of 10 nM of each fluorescently labeled readout probe (Cy5 or IR800) in a 10% hybridization buffer. Unbound readout probes were then removed by washing the sample with a 10% formamide wash buffer, followed by a wash with 2× SSC. Next, we flowed the imaging buffer into the chamber before image acquisition. The imaging buffer consisted of 50 mM Tris-HCl pH 8, 2 mM Trolox (Sigma, cat. no. 238813), 10% glucose, 0.5 mg/ml glucose oxidase (Sigma, cat. no. G2133) and 40 μg/ml catalase (Sigma, cat. no. C30-100MG) in 2x SSC. Fluorescent signals were removed by flowing 2x SSC into the chamber and photobleaching the sample by exposure to 647 nm and 750 nm lasers for 20 s. This hybridization and photobleaching cycle was repeated until all the bits were imaged. The readout probes for Guidestar were imaged first, followed by the MERFISH readouts, to ensure optimal quality of the Guidestar images and avoid potential signal degradation arising from long acquisition times. The total number of hybridization rounds (each round consists of 2 color channels) is the sum of 4 Guidestar rounds (8 Guidestar genes) and 8 MERFISH rounds (16 bits MHD4 MERFISH).

### Data Analysis for RNA decoding

#### Image preprocessing and MERFISH analysis

We used a modified version of the analysis pipeline described previously^23^ to identify RNA spots. Pre-processing steps (image registration, image filtering, normalization) and comparison to codewords were performed as previously described. For the pre-processing parameters, we set the minimum magnitude threshold (mean intensity) per spot to 0.1, the maximum distance threshold (Euclidean distance to a unit-normalized gene’s codeword) to 0.605 and the minimum spot size (pixel region) to 2 pixels. Bit normalization and image filtering parameters were used as previously described.

#### Implementation of blank fraction thresholding

We further implemented blank fraction thresholding (BFT) to identify RNA spots at target misidentification rate following the approach described by Xia et al.^19^. Briefly, a 3D histogram of mean intensity, minimum distance to gene’s code-words and number of spots was constructed. A similar 3D histogram was constructed for seven blank barcodes. For each histogram bin, we calculated the fraction of blank barcodes over total gene + blank barcodes as the blank fraction score (BFS). A high BFS suggests a high probability of barcode misidentification for that histogram bin. We then set a threshold to filter spots by their BFS, which is determined by the histogram bin matching each spots’ characteristics. This threshold is tuned to achieve the desired misidentification rate which is defined as mean count per blank barcode / mean count per gene barcode. The gross misidentification rate of all barcodes is conventionally set as 0.05, which can be interpreted as a 5% chance that any spot called (after filtering) would be an artifactual spot rather than a true RNA molecule.

#### Signal and background brightness comparison

To quantify the improvement in Guidestar bits’ image quality over that of MERFISH bits, we randomly sampled 500 spots in Guidestar and MERFISH images. The center of each spot was estimated by detecting local maxima and the pixel intensity was plotted as a function of distance to the spot center. The central pixel intensity was taken to be the peak signal while the furthest pixels’ mean intensity within a 10x10 ROI was taken to be the background noise. Signal-to-background ratio was calculated as the mean intensity of pixels at each distance divided by the background intensity (**Figure S1b**).

#### Colocalization analysis

We used the *peaklocalmax* function of the *Scikit-image* v0.19.3 library^22^ in Python to find Guidestar spots. The spots were detected by finding local maxima in the filtered image. Only peaks above an intensity threshold were called out as Guidestar spots. We set the intensity threshold for each of the Guidestar genes through visual inspection of the images. To set a maximum distance between spots below which they are considered colocalized, we considered a range of colocalization distance thresholds (**Figure S2a, c**) and selected a euclidean distance of 4 pixels or 480nm. A MERFISH callout is considered colocalized if there is a Guidestar spot of the corresponding gene within a euclidean distance of 4 pixels from the centroid of the MERFISH spot.

### Training of classifiers using Guidestar

#### Quality control and Guidestar gene selection

Some probe sets may have image quality issues due to nonspecific binding, poor probe penetrance, differences in probe binding kinetics, or blocking by RNA-binding proteins. Hence, we performed a quality control step on each of the 7 Guidestar genes in the library to ensure that the Guidestar genes used yielded high quality images (**Figure S1a**). We removed *Igf1*, *Pigr* and *Cps1* for the AML-12 cell line sample as these RNAs have low FPKM (the probe-library was designed for liver tissue data). We then tested all possible combinations of the remaining 4 Guidestar genes using 5-fold cross validation and a random forest classifier, and found that using all of the 4 high-quality genes: *Acly*, *Gpam*, *Hnf4a* and *Ube2z* yielded the highest F1 score (**Figure S3a**). After optimizing gene combinations, we retrained the model using these 4 genes on the full training set and evaluated its performance on the held-out test set.

#### Class balancing approach

As we observed varying numbers of positive training examples (spots colocalized between guidestar and MERFISH) and negative training examples (spots found in MERFISH only), we tested a variety of approaches to even out class imbalances (**Figure S3c, d**). We sought to use as many of the colocalized spots as possible as they provide valuable information for training the classifier. Therefore, if there were more MERFISH-only than colocalized spots, we downsampled MERFISH-only spots randomly to attain a 1:1 ratio with colocalized spots within each field of view (FOV). On the other hand, if there were more colocalized than MERFISH-only spots, we did not downsample the colocalized spots. Instead, we augmented our negative examples by adding spots corresponding to blank code-words. To prevent the model from learning excessively from blanks, we did not add more blanks than MERFISH-only spots, such that blanks and MERFISH-only spots were at a 1:1 ratio.

#### Additional features and feature selection

Conventionally, the mean intensity of pixels within the spot, the hamming distance between the imaged codeword and the closest theoretical barcode, and the number of pixels in the spot (spot size) are considered in calculating the blank fraction score. We expanded on these 3 features to include intensity and distance-related characteristics of second and third closest theoretical barcodes **(Table S3**). We then performed sequential forward feature selection to obtain the feature set (1. mean intensity of pixels in the on-bits of the closest theoretical codeword; 2. smallest distance between imaged codeword and second closest theoretical codeword; 3. smallest distance between imaged codeword and third closest theoretical codeword; 4. Number of pixels in the spot) that produces the best F1 score during cross validation out of the 15 features we extracted from the data (**Figure S3e**).

#### Model training

We aimed to construct a classifier for binary classification (0 = not RNA callout, 1 = true RNA callout). We then labeled all putative MERFISH callouts (without BFT) as the following: MERFISH callouts that colocalized with Guidestar spots were labeled as 1 (gene spot). Whereas, MERFISH callouts that did not colocalize with Guidestar spots were labeled as 0 (not gene spot). We used 80% of the data as the training set while the remaining 20% served as the test set. We tested all possible combinations of Guidestar genes (**Figure S3a, b**) and sampling methods to reduce class imbalance (**Figure S3c, d**). After optimizing for combinations and numbers of genes, we found that using a smaller subset of genes did not improve accuracy. For this reason, and to maximize the diversity of the training data and improve generalizability, we used all Guidestar genes that passed initial QC. For the cell line dataset, we found that the best sampling method was to augment the minority negative class with spots decoded as blanks instead of downsampling the majority positive class which contained valuable information on colocalized spots. For the tissue sample dataset, we downsampled the majority negative class as there were no sources for positive class augmentation. We performed sequential forward feature selection using SequentialFeatureSelector from the *mlxtend* v0.22.0^30^ library in Python (**Figure S3e**, **Table S4**) to optimize the features used for both datasets separately. Finally, we optimized hyperparameters of the *RandomForestClassifier* from the *sklearn* v1.1.1^31^ library in Python (**Figure S3f**). The optimized hyperparameter set used was: min_samples_split=0.001, min_impurity_decrease=0.01, max_features=1, max_depth=12, criterion=’gini’, min_weight_fraction_leaf=0.01, n_estimators=100, max_samples=0.8. For comparison to the BFT method, we used the default output probability threshold of the *RandomForestClassifier.predict* function. All model optimization was conducted via 5-fold cross validation on the training set to maximize the F1 score.

#### Classifier model and BFT evaluation

To determine precision and recall, we began with candidate RNA callouts generated by the decoding pipeline prior to spot filtering^23^. The pipeline was first run with relaxed thresholds (greater than 0.1 mean intensity, within 0.605 distance to closest codeword, at least 2 pixels in size) to identify a maximal set of putative RNA spots. These putative RNA spots were divided into two groups: those co-localizing with Guidestar spots, considered “true” RNA spots, and those without co-localization, defined as “false” RNA spots. Precision was defined as the number of “true” RNA spots that are called out by the model, divided by the total number of spots called by the model. Recall was defined as the number of “true” RNA spots that are called out by the model, divided by the total number of “true” RNA spots. The F1 score was computed as 2 * precision * recall / (precision + recall). See **Figure 2a** for more details. The ‘true’ RNA spots are the intersections of both putative MERFISH and Guidestar spots, indicating that they are high-confidence ground truth examples for training and validation.

The optimized model was applied to the test set and the precision, recall, and F1 score for each FOV was calculated. We also generated the receiver operating characteristic curve (ROC) for each FOV in the test set (**Figure 2d**, dashed lines) and reported the mean area under the curve across FOVs. The true positive rates of individual FOV’s ROC (**Figure 2d**, **Figure S4a**, dashed lines) were interpolated across false positive rates ranging from 0 to 1, and averaged to obtain the mean ROC curve (**Figure 2d**, **Figure S4b**, solid lines). The optimized model was applied to all other genes in the library. We calculated Pearson’s correlation of log count values between bulk FPKM and gene spot counts for each FOV as well as over the full dataset. The FPKM values from bulk RNA-seq of mouse liver tissues were downloaded from the ENCODE portal (<u>ENCFF844MJF</u> and <u>ENCFF271DWG</u>) and averaged. **Figure S5** shows the correlation of BFT/Guidestar model callouts with the bulk RNA-seq FPKM.

