## Supplementary Figures and Tables for "Guidestar: a spike-in approach to improve RNA detection accuracy in imaging-based spatial transcriptomics"

### Supplemental Information

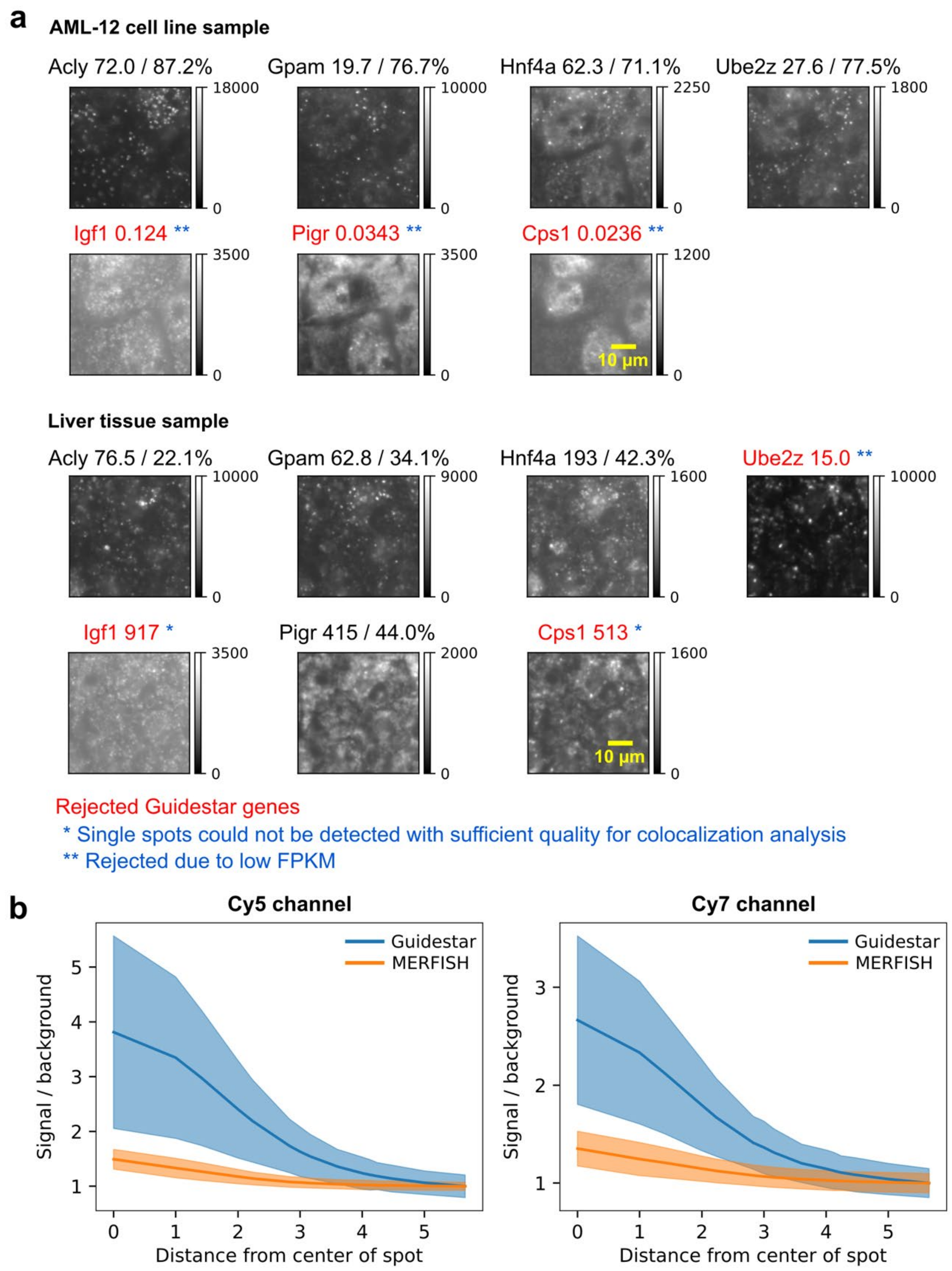

**Figure S1: (a)** Representative raw images from Guidestar imaging rounds. Genes with good image quality and distinctive spots were selected and used for training the random forest classifier, while rejected genes are indicated in red. Numbers next to the gene names indicate FPKM, while the percentages indicate colocalization percentages. Genes were rejected due low FPKM (**) or when distinct spots could not be detected with sufficient quality for colocalization analysis (*). **(b)** Fold change in signal intensity (at each distance) over background noise (mean pixel intensity at the edges of a 10x10 ROI from spot center) in cell line data. The steeper curve for Guidestar spots indicates more distinction of signal from background in Guidestar images and better resolved point spread function. Lines indicate the mean while the shaded region indicates the standard deviation.

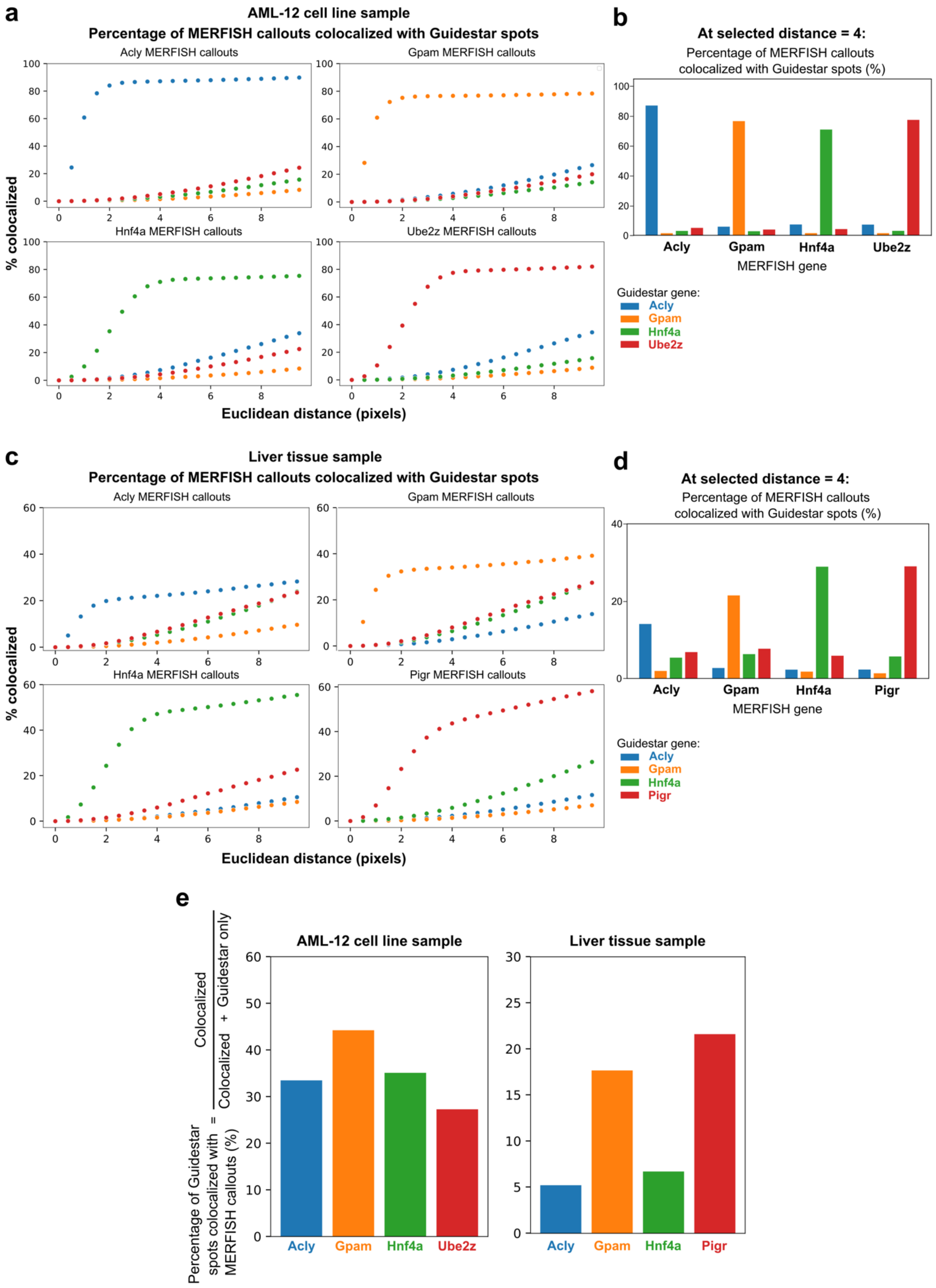
a
AML-12 cell line sample
Percentage of MERFISH callouts colocalized with Guidestar spots
Acly MERFISH callouts
Gpam MERFISH callouts
Hnf4a MERFISH callouts
Ube2z MERFISH callouts
% colocalized
Euclidean distance (pixels)
b
At selected distance = 4:
Percentage of MERFISH callouts colocalized with Guidestar spots (%)
Acly
Gpam
Hnf4a
Ube2z
MERFISH gene
Guidestar gene:
Acly
Gpam
Hnf4a
Ube2z
c
Liver tissue sample
Percentage of MERFISH callouts colocalized with Guidestar spots
Acly MERFISH callouts
Gpam MERFISH callouts
Hnf4a MERFISH callouts
Pigr MERFISH callouts
% colocalized
Euclidean distance (pixels)
d
At selected distance = 4:
Percentage of MERFISH callouts colocalized with Guidestar spots (%)
Acly
Gpam
Hnf4a
Pigr
MERFISH gene
Guidestar gene:
Acly
Gpam
Hnf4a
Pigr
e
Percentage of Guidestar spots colocalized with MERFISH callouts (%) = Colocalized / (Colocalized + Guidestar only)
AML-12 cell line sample
Liver tissue sample
Acly
Gpam
Hnf4a
Ube2z
Acly
Gpam
Hnf4a
Pigr

**Figure S2:** Tuning of colocalization distance for **(a)** AML-12 cell line sample and **(c)** liver tissue sample. The distance threshold at which the colocalization percentage plateaus was selected as the optimal threshold (4 pixels ~ 0.48um), above which colocalization becomes non-specific. The percentage of MERFISH spots colocalized with Guidestar spots (bars) for each Guidestar gene for **(b)** AML-12 cell line sample and **(d)** liver tissue sample, at the optimal threshold. There is high percentage of colocalization between corresponding MERFISH and Guidestar genes (71.1% - 87.2%) while there is low percentage of colocalization between unmatched MERFISH and Guidestar genes (1.54% - 7.41%), indicating high specificity of colocalization. For the liver tissue, while the challenging sample has poorer colocalization, the percentage of colocalization between corresponding MERFISH and Guidestar genes (22.1% - 44.0%) is greater than that between unmatched MERFISH and Guidestar genes (1.34% - 8.54%). **(e)** The percentage of Guidestar spots colocalized with MERFISH spots (bars) for each Guidestar gene (left: AML-12 cell line sample; right: liver tissue sample).

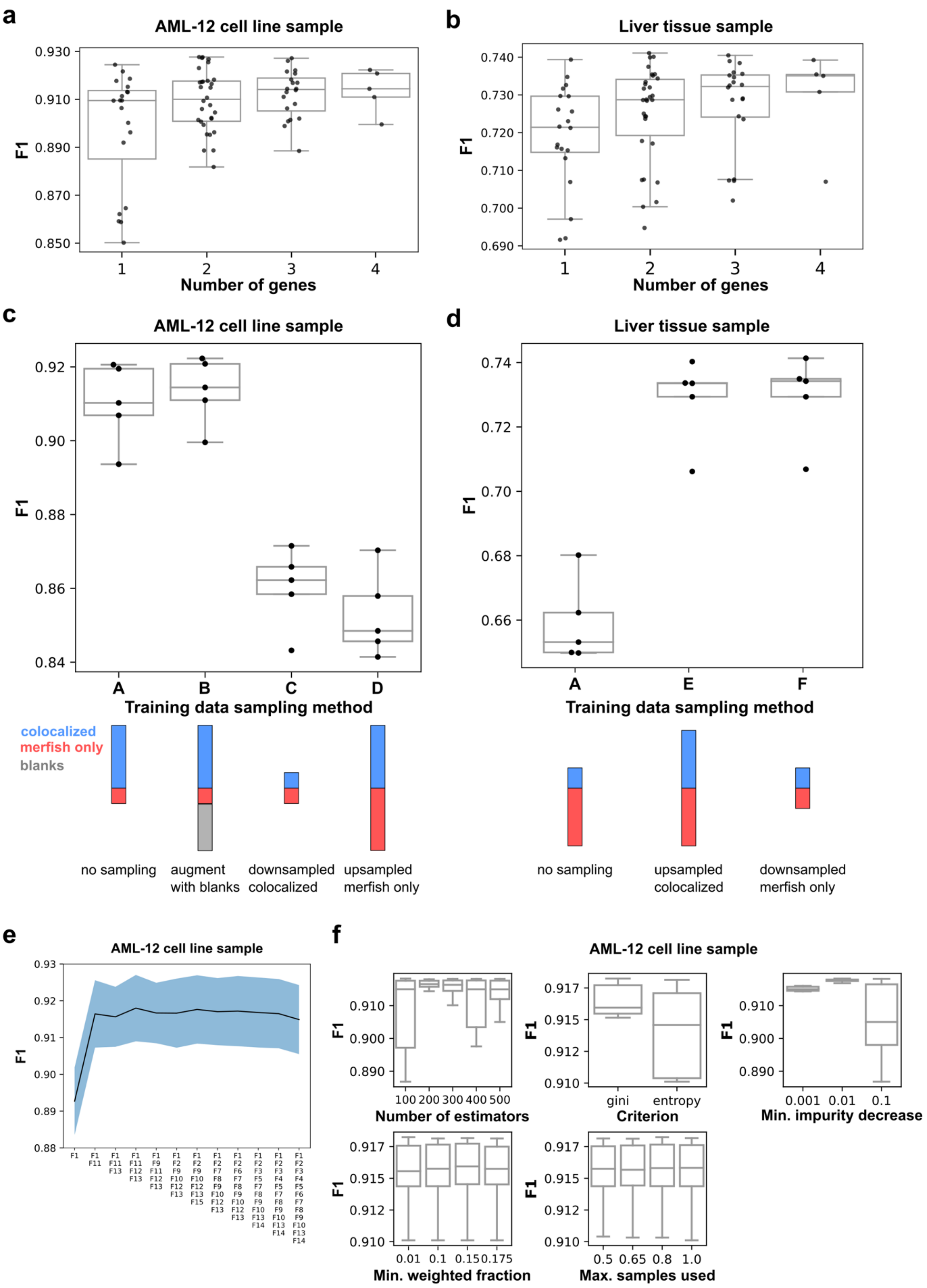


**Figure S3:** Effects of increasing the number of Guidestar genes used on F1 score (5-fold cross validation) for **(a)** AML-12 cell line sample and **(b)** liver tissue sample. Effect of sampling (class balancing)

methods on F1 score (5-fold cross validation). **(c) AML-12 cell line sample:** A: All colocalized spots and all MERFISH only spots; B: Augment minority class (MERFISH only spots) with spots decoded as blanks such that positive and negative classes are equal; C: Downsampling of majority class (colocalized spots) such that positive and negative classes are equal; D: Upsampling of minority class such that positive and negative classes are equal. **(d) Liver tissue sample:** A: No sampling; E: Upsampling of minority class (colocalized spots) such that positive and negative classes are equal; F: Downsampling of majority class (MERFISH only spots) such that positive and negative classes are equal. Positive class refers to colocalized spots while negative class refers to MERFISH only spots. **(e)** Sequential forward floating feature selection was performed via 5-fold cross validation to maximize the F1 score for the AML-12 murine liver cell sample. As performance stopped improving beyond 4-5 features, feature selection was stopped at 12 features. The black line shows the mean while the shaded blue area indicates the standard deviation across cross validation sets. Details of features are described in **Table S4**. **(f)** Hyperparameter optimization of random forest classifier to maximize F1 score via 5-fold cross validation. The hyperparameters optimized were the number of estimators in the random forest, criterion for leaf node impurity calculation, the minimum decrease in impurity for a leaf node to be split, the minimum weighted fraction of the sum of total sample weights at a leaf node and the maximum proportion of samples used to train each estimator in the random forest. See methods for hyperparameters used in the model. Center line, median; height of the box, interquartile range (IQR); whiskers, 1.5 × IQR.

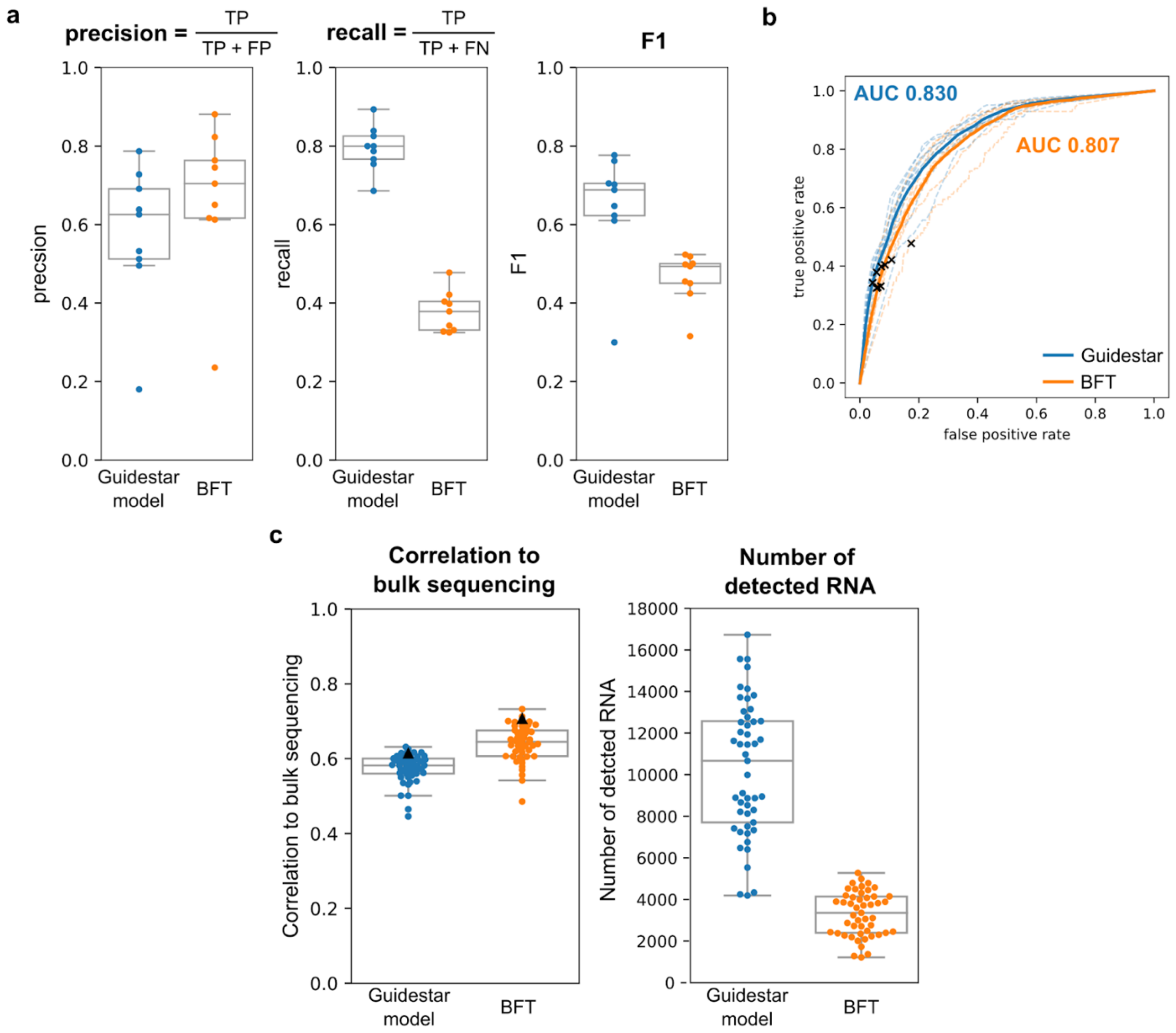

**Figure S4:** The Guidestar model optimized on the AML-12 cell line sample can be retrained on a liver tissue dataset to yield improvements over the BFT. (a) The Guidestar model has reduced precision but increased recall, leading to an overall increase in F1 compared to BFT at 0.05 misidentification rate in the test set. Each point represents a different FOV, which serves as technical replicates, in the test set. (b) The Guidestar model has a similar average receiver-operator-curve (solid lines) to the BFT with slightly higher area-under-the-curve (AUC). BFT performance at 0.05 misidentification rate is indicated by the crosses for each FOV in the test set (dashed lines). AUC values are indicated for the average ROC. (c) When applied to the full library of genes, the Guidestar model filtered counts achieves lower correlation to bulk sequencing but detects 3.1x more RNA than BFT. Center line, median; height of the box, interquartile range (IQR); whiskers, 1.5 × IQR.

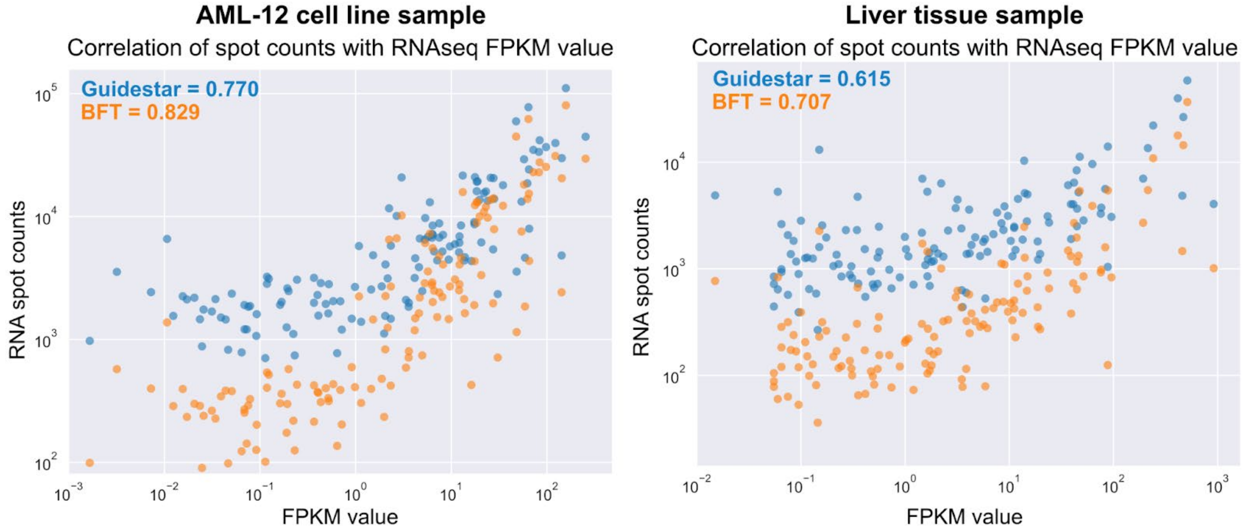


**Figure S5:** The correlation between bulk RNA sequencing RNA FPKM and BFT (orange) or Guidestar model (blue) for AML-12 cell line (left) and liver tissue sample (right).

**Table S2: 133-genes Guidestar library readout sequences**

| *Bit* | *Readout Sequence* |
|---|---|
| B1 | /5IRD800CWN/CGCAACGCTTGGGACGGTTCCAATCGGATC |
| B2 | /5IRD800CWN/CGAATGCTCTGGCCTCGAACGAACGATAGC |
| B3 | /5IRD800CWN/ACAAATCCGACCAGATCGGACGATCATGGG |
| B4 | /5IRD800CWN/CAAGTATGCAGCGCGATTGACCGTCTCGTT |
| B5 | /5IRD800CWN/GCGGGAAGCACGTGGATTAGGGCATCGACC |
| B6 | /5IRD800CWN/AAGTCGTACGCCGATGCGCAGCAATTCACT |
| B7 | /5IRD800CWN/CGAAACATCGGCCACGGTCCCGTTGAACTT |
| B8 | /5IRD800CWN/ACGAATCCACCGTCCAGCGCGTCAAACAGA |
| B9 | CGCGAAATCCCCGTAACGAGCGTCCCTTGC/3Cy5Sp/ |
| B10 | GCATGAGTTGCCTGGCGTTGCGACGACTAA/3Cy5Sp/ |
| B11 | CCGTCGTCTCCGGTCCACCGTTGCGCTTAC/3Cy5Sp/ |
| B12 | GGCCAATGGCCCAGGTCCGTCACGCAATTT/3Cy5Sp/ |
| B13 | TTGATCGAATCGGAGCGTAGCGGAATCTGC/3Cy5Sp/ |
| B14 | CGCGCGGATCCGCTTGTCGGGAACGGATAC/3Cy5Sp/ |
| B15 | GCCTCGATTACGACGGATGTAATTCGGCCG/3Cy5Sp/ |
| B16 | GCCCGTATTCCCGCTTGCGAGTAGGGCAAT/3Cy5Sp/ |
| B17 | /5IRD800CWN/GTGTGGTGGTTAATATATGTAAGTGGGTGG |
| B18 | /5IRD800CWN/GAAGGGAGATGGGATTTAGGGAGGTGGGTG |
| B19 | /5IRD800CWN/TGATGAAGGGAGGAAGTTAGGATGGGTTGT |
| B20 | /5IRD800CWN/GTATGTGGGTATGGAGAAGGGAGTAATTGA |
| B21 | TGGATGGGTGGAGATGTTGTGAAGAAATAG/3Cy5Sp/ |
| B22 | AATGGTGAGAGGAAGGTGATGTAGTGGGAT/3Cy5Sp/ |
| B23 | ATTATGTAGGATGTGGAAGGAGTAGAGGAG/3Cy5Sp/ |

**Table S3: Multiplexed FISH library codebook**

| *Gene* | *Code-word* |
|---|---|
| Igf1 | 0 0 1 0 0 0 0 0 0 1 1 0 0 0 1 0 |
| Cps1 | 1 0 0 0 0 0 0 1 1 0 0 0 1 0 0 0 |
| Glul | 0 0 0 0 1 1 0 1 0 0 0 1 0 0 0 0 |
| Glud1 | 0 0 0 0 0 0 0 1 0 0 0 0 1 1 1 0 |
| Pigr | 0 1 0 0 0 0 1 0 0 0 0 1 0 0 0 1 |
| Pck1 | 1 0 0 1 0 0 0 0 0 0 1 1 0 0 0 0 |
| Uox | 0 1 0 0 0 0 0 0 1 1 0 0 0 1 0 0 |
| Hnf4a | 0 0 0 0 1 1 0 0 1 0 0 0 0 0 0 1 |
| Abcc3 | 0 1 0 1 0 0 0 0 0 1 0 0 0 0 0 1 |
| G6pc | 0 0 1 0 1 0 1 1 0 0 0 0 0 0 0 0 |
| Paqr9 | 0 0 0 0 0 1 1 0 0 1 0 0 0 0 0 1 |
| Cpt1a | 0 1 0 0 1 0 0 1 0 1 0 0 0 0 0 0 |
| Acly | 1 0 0 0 0 0 1 0 0 1 1 0 0 0 0 0 |
| Ganab | 0 0 0 1 0 0 0 0 1 1 1 0 0 0 0 0 |
| Gpam | 0 0 1 0 1 0 0 0 0 1 0 0 0 1 0 0 |
| Plxnb2 | 0 0 1 0 0 1 0 1 0 0 0 0 0 0 1 0 |
| Dpyd | 0 1 1 1 0 0 0 0 0 0 0 1 0 0 0 0 |
| Clptm1 | 0 0 0 1 0 0 1 0 0 1 0 1 0 0 0 0 |
| Ddx17 | 0 1 0 1 0 1 1 0 0 0 0 0 0 0 0 0 |
| Scap | 0 0 1 0 0 0 0 0 0 0 0 0 0 1 1 1 |
| Uba1 | 0 0 0 1 1 0 1 0 0 0 1 0 0 0 0 0 |
| Gns | 0 1 0 1 1 0 0 0 0 0 0 0 0 0 1 0 |
| Lrrc3 | 0 1 0 0 0 0 1 1 1 0 0 0 0 0 0 0 |
| Fads6 | 0 1 0 0 0 0 0 0 1 0 1 0 0 0 0 1 |
| Hs3st3b1 | 1 0 1 0 0 0 1 0 0 0 0 0 0 0 1 0 |
| Zcchc24 | 0 1 0 0 0 0 1 0 0 0 1 0 0 0 1 0 |
| Cdc42bpb | 1 0 0 0 1 0 0 0 0 0 0 0 1 1 0 0 |
| Zdhhc5 | 1 0 1 0 1 1 0 0 0 0 0 0 0 0 0 0 |
| Pdxk | 0 0 1 0 0 1 0 0 1 0 1 0 0 0 0 0 |
| Vwa8 | 1 0 0 0 0 0 1 0 0 0 0 0 0 1 0 1 |
| Flii | 0 0 1 0 0 0 0 0 0 1 0 1 0 0 0 1 |
| Ube2z | 0 0 0 0 0 0 1 1 0 0 0 1 0 0 1 0 |
| Slc38a2 | 0 0 0 0 0 0 1 1 0 1 0 0 0 1 0 0 |
| Stat2 | 0 1 0 0 0 0 0 1 0 0 1 1 0 0 0 0 |
| Bcl9l | 0 0 0 0 1 0 0 0 0 1 1 1 0 0 0 0 |
| Tomm20 | 1 0 1 1 0 0 0 0 0 0 0 0 0 1 0 0 |
| Il4ra | 0 0 0 0 1 1 1 0 0 0 0 0 0 0 1 0 |
| Nol6 | 0 1 0 0 0 0 0 0 1 0 0 1 0 0 1 0 |
| Ide | 1 1 0 0 0 0 1 0 0 0 0 0 1 0 0 0 |
| Ddx3x | 0 0 0 0 0 0 0 1 1 0 1 0 0 0 1 0 |
| Rnf144b | 1 1 0 0 0 1 0 0 1 0 0 0 0 0 0 0 |
| Mybbp1a | 1 1 0 0 0 0 0 0 0 0 0 1 0 1 0 0 |
| Hnf1b | 1 0 0 0 1 0 0 0 1 0 1 0 0 0 0 0 |
| Flnb | 0 1 0 1 0 0 0 0 0 0 1 0 0 1 0 0 |
| Golga2 | 0 0 0 0 0 0 0 0 0 1 1 0 0 1 0 1 |
| Cdh5 | 0 0 1 1 1 0 0 0 0 0 0 0 1 0 0 0 |
| Tbx3 | 0 0 0 0 1 0 0 0 0 0 1 0 0 1 1 0 |
| Paxip1 | 0 0 1 0 0 0 1 0 0 0 1 0 1 0 0 0 |
| Dennd5a | 1 0 0 0 0 0 0 0 1 1 0 1 0 0 0 0 |

| | |
|---|---|
| Rhbdd2 | 0 0 0 0 0 1 0 1 0 0 0 0 0 1 0 1 |
| Mob3b | 0 0 1 0 0 1 0 0 0 0 0 0 1 1 0 0 |
| Plekhm2 | 1 0 0 0 0 0 0 0 0 0 1 0 1 0 1 0 |
| Ube3a | 0 0 0 1 1 1 0 0 0 0 0 0 0 1 0 0 |
| Dgkq | 1 0 0 0 0 1 1 1 0 0 0 0 0 0 0 0 |
| Tecpr1 | 0 0 0 1 0 0 1 0 0 0 0 0 0 1 1 0 |
| Iffo2 | 1 0 0 1 0 0 1 0 1 0 0 0 0 0 0 0 |
| Zhx2 | 0 0 0 0 1 0 1 0 1 1 0 0 0 0 0 0 |
| Gpd2 | 0 1 1 0 0 1 0 0 0 0 0 0 0 0 0 1 |
| Lrat | 0 0 0 1 0 0 0 1 0 1 0 0 0 0 1 0 |
| Lrtm1 | 1 0 0 1 0 1 0 0 0 0 0 0 1 0 0 0 |
| Tbl1xr1 | 0 1 0 0 0 0 0 0 0 0 0 0 1 1 0 1 |
| Pecam1 | 0 0 0 0 1 0 0 0 1 0 0 0 1 0 1 0 |
| Cog3 | 0 0 0 0 0 0 1 0 1 0 1 1 0 0 0 0 |
| Atp11b | 0 0 1 0 0 1 1 0 0 0 0 1 0 0 0 0 |
| Add3 | 0 1 1 0 1 0 0 0 1 0 0 0 0 0 0 0 |
| Mcam | 0 1 0 1 0 0 0 0 1 0 0 0 1 0 0 0 |
| 1700017B05Rik | 0 1 0 0 1 0 1 0 0 0 0 0 0 1 0 0 |
| Megf8 | 1 0 1 0 0 0 0 1 0 0 0 1 0 0 0 0 |
| Tppp | 1 0 0 0 0 0 0 0 1 0 0 0 0 1 1 0 |
| Vps13c | 1 1 0 0 1 0 0 0 0 0 0 0 0 0 0 1 |
| Exoc2 | 0 0 1 0 0 0 0 0 1 0 0 1 1 0 0 0 |
| Taok1 | 1 0 0 0 0 1 0 0 0 1 0 0 0 1 0 0 |
| Enpp4 | 0 0 0 0 0 1 1 0 0 0 1 0 0 1 0 0 |
| Des | 0 0 0 0 0 0 1 0 0 1 0 0 1 0 1 0 |
| Ranbp2 | 0 0 0 1 1 0 0 1 0 0 0 0 0 0 0 1 |
| Golga4 | 1 0 0 0 1 0 0 1 0 0 0 0 0 0 1 0 |
| Sptlc2 | 0 0 0 0 0 0 0 1 1 0 0 1 0 0 0 1 |
| Sox9 | 0 0 1 0 0 0 1 0 1 0 0 0 0 1 0 0 |
| Capn5 | 0 0 0 1 1 0 0 0 1 0 0 1 0 0 0 0 |
| Rasa1 | 0 0 0 0 0 0 1 0 1 0 0 0 0 0 1 1 |
| Vcpip1 | 1 0 1 0 0 0 0 0 0 1 0 0 1 0 0 0 |
| Tln2 | 1 0 0 1 1 0 0 0 0 1 0 0 0 0 0 0 |
| Gpc1 | 0 0 0 0 0 0 1 1 0 0 1 0 0 0 0 1 |
| Adcy7 | 0 1 0 0 0 1 0 1 0 0 0 0 1 0 0 0 |
| Zxdb | 0 0 0 0 0 1 0 0 1 1 0 0 0 0 1 0 |
| Cdk19 | 0 0 0 1 0 0 0 0 1 0 0 0 0 1 0 1 |
| Chic1 | 0 1 0 0 0 1 0 0 0 1 0 1 0 0 0 0 |
| C3ar1 | 0 0 1 0 0 0 0 0 0 0 1 1 0 1 0 0 |
| Ccdc88b | 0 0 1 1 0 1 0 0 0 1 0 0 0 0 0 0 |
| Aatk | 0 0 0 1 0 1 0 0 0 0 0 1 0 0 0 1 |
| Tmed8 | 1 0 0 0 0 0 0 1 0 1 0 0 0 0 0 1 |
| Zfp174 | 1 0 1 0 0 0 0 0 1 0 0 0 0 0 0 1 |
| Fastkd2 | 0 0 0 0 0 1 0 1 0 1 1 0 0 0 0 0 |
| Zfp870 | 0 0 0 1 0 0 0 0 0 0 1 0 1 0 0 1 |
| Dsg1c | 0 0 0 0 1 0 0 0 0 1 0 0 0 0 1 1 |
| Wdr19 | 0 0 0 0 1 1 0 0 0 1 0 0 1 0 0 0 |
| Mphosph9 | 0 0 1 0 1 0 0 0 0 0 0 1 0 0 1 0 |
| Abca5 | 0 0 0 0 0 0 0 0 1 0 1 0 1 1 0 0 |
| Atp8b5 | 0 0 0 1 0 0 0 1 0 0 0 1 0 1 0 0 |
| Ankrd26 | 0 0 1 0 0 0 0 1 0 0 0 0 1 0 0 1 |

| | |
|---|---|
| Tbc1d9 | 0 1 1 0 0 0 0 0 0 0 0 0 0 1 0 1 0 |
| Zfp560 | 0 0 1 0 1 0 0 0 0 0 0 1 0 0 0 0 1 |
| Ggt1 | 0 1 0 0 0 1 0 0 0 0 0 0 0 0 1 1 0 |
| AW554918 | 0 1 0 0 0 0 0 0 0 0 1 1 0 1 0 0 0 |
| Itga4 | 0 0 1 1 0 0 0 1 0 0 1 0 0 0 0 0 0 |
| Ammecr1 | 0 0 0 0 0 1 0 0 1 0 0 1 0 1 0 0 0 |
| Hecw2 | 0 0 0 0 0 0 0 0 0 0 0 1 1 0 0 1 1 |
| Fam102b | 0 0 0 0 1 0 0 0 0 0 0 0 1 0 1 0 1 |
| Nova2 | 0 0 0 0 0 0 0 0 0 0 1 0 1 0 1 1 0 |
| Krt7 | 1 0 0 0 1 0 1 0 0 0 0 1 0 0 0 0 0 |
| Prrg3 | 0 0 0 0 0 1 0 0 0 0 0 0 0 1 0 1 1 |
| Kif7 | 0 0 0 1 0 0 0 0 0 0 0 0 1 1 0 1 0 |
| Setbp1 | 1 0 0 0 0 0 0 1 0 0 1 0 0 1 0 0 0 |
| Pcdh17 | 0 0 1 1 0 0 0 0 1 0 0 0 0 0 0 1 0 |
| Brip1 | 0 0 0 1 0 0 1 1 0 0 0 0 1 0 0 0 0 |
| Armcx4 | 0 0 0 0 0 1 1 0 1 0 0 0 1 0 0 0 0 |
| Scube1 | 0 0 0 0 1 0 0 1 1 0 0 0 0 1 0 0 0 |
| Alms1 | 0 0 0 0 0 0 0 0 1 1 0 0 1 0 0 1 0 |
| Dync2h1 | 1 0 0 1 0 0 0 0 0 0 0 0 0 0 0 1 1 |
| Hcn3 | 1 1 0 1 0 0 0 1 0 0 0 0 0 0 0 0 0 |
| Sdk1 | 0 1 1 0 0 0 0 1 0 0 0 0 0 1 0 0 0 |
| Slit3 | 0 0 0 0 0 0 0 1 0 1 0 1 1 0 0 0 0 |
| Plag1 | 1 0 0 0 0 1 0 0 0 0 0 1 0 0 1 0 0 |
| Sema3g | 0 0 0 0 0 0 1 0 0 0 0 1 1 1 0 0 0 |
| Ncs1 | 0 0 0 1 0 0 0 0 0 1 0 0 1 1 0 0 0 |
| Egflam | 0 1 0 0 1 1 0 0 0 0 1 0 0 0 0 0 0 |
| Lama3 | 1 0 0 0 0 0 0 0 0 0 0 0 1 1 0 0 1 |
| Igdcc4 | 0 0 0 0 0 1 0 0 0 0 1 1 1 0 0 0 0 |
| Sgpp2 | 0 0 0 0 1 0 0 1 0 0 1 0 1 0 0 0 0 |
| Lrrc16b | 0 0 0 1 0 1 0 0 0 0 1 0 0 0 0 1 0 |
| Col4a4 | 1 0 0 0 0 1 0 0 0 0 1 0 0 0 0 0 1 |
| Cnnm1 | 0 1 0 0 0 0 0 1 0 0 0 0 0 0 0 1 1 |
| Cftr | 0 1 1 0 0 0 1 0 0 1 0 0 0 0 0 0 0 |
| Blank1 | 0 0 0 1 0 1 0 1 1 0 0 0 0 0 0 0 0 |
| Blank2 | 0 0 1 0 0 0 0 1 1 1 0 0 0 0 0 0 0 |
| Blank3 | 1 1 1 0 0 0 0 0 0 0 1 0 0 0 0 0 0 |
| Blank4 | 0 1 0 0 1 0 0 0 0 0 0 1 1 0 0 0 0 |
| Blank5 | 0 0 1 1 0 0 1 0 0 0 0 0 0 0 0 0 1 |
| Blank6 | 1 1 0 0 0 0 0 0 0 1 0 0 0 0 0 1 0 |
| Blank7 | 0 0 0 0 1 0 1 0 0 0 0 0 0 1 0 0 1 |

| | Feature |
|---|---|
| **F1** | **Mean intensity of pixels in the on-bits of the closest theoretical barcode in spot** |
| F2 | Mean intensity of pixels in the off-bits of the closest theoretical barcode in spot |
| F3 | Mean intensity of pixels in the on-bits of the second closest theoretical barcode in spot |
| F4 | Mean intensity of pixels in the off-bits of the second closest theoretical barcode in spot |
| F5 | Mean intensity of pixels in the on-bits of the third closest theoretical barcode in spot |
| F6 | Mean intensity of pixels in the off-bits of the third closest theoretical barcode in spot |
| F7 | Ratio of F1 to F3 |
| F8 | Ratio of F1 to F5 |
| F9 | Ratio of F1 to F2 |
| **F10** | **Smallest distance between imaged spot codeword and the closest theoretical barcode** |
| F11 | Smallest distance between imaged spot codeword and the second closest theoretical barcode |
| F12 | Smallest distance between imaged spot codeword and the third closest theoretical barcode |
| **F13** | **Number of pixels in the spot** |
| F14 | Ratio of the long edge to the short edge of the spot |
| F15 | Number of disconnected pixel clusters within the spot |

**Table S4:** Descriptions of spot features used in Random Forest Classifier. Conventional features are highlighted.
